# Quantifying the impact of Human Leukocyte Antigen on the human gut microbiome

**DOI:** 10.1101/2020.01.14.907196

**Authors:** Stijn P. Andeweg, Can Keşmir, Bas E. Dutilh

**Affiliations:** Theoretical Biology and Bioinformatics, Science for Life, Utrecht University, Padualaan 8, 3584 CH, Utrecht, the Netherlands

## Abstract

**Objective:** The gut microbiome is affected by a number of factors, including the innate and adaptive immune system. The major histocompatibility complex (MHC), or the human leukocyte antigen (HLA) in humans, performs an essential role in vertebrate immunity, and is very polymorphic in different populations. HLA determines the specificity of T lymphocyte and natural killer (NK) cell responses, including against the commensal bacteria present in the human gut. Thus, it is likely that our HLA molecules and thereby the adaptive immune response, can shape the composition of our microbiome. Here, we investigated the effect of HLA haplotype on the microbiome composition.

**Results:** We performed HLA typing and microbiota composition analyses on 3,002 public human gut microbiome datasets. We found that (i) individuals with functionally similar HLA molecules (i.e. presenting similar peptides) are also similar in their microbiota, and (ii) HLA homozygosity correlated with microbiome diversity, suggesting that diverse immune responses limit microbiome diversity.

**Conclusion:** Our results show a statistical association between host HLA haplotype and gut microbiome composition. Because the HLA haplotype is a readily measurable parameter of the human immune system, these results open the door to incorporating the immune system into predictive microbiome models.

**IMPORTANCE:** The microorganisms that live in the digestive tracts of humans, known as the gut microbiome, are essential for hosts survival as they support crucial functions. For example, they support the host in facilitating the uptake of nutrients and give colonization resistance against pathogens. The composition of the gut microbiome varies among humans. Studies have proposed multiple factors driving the observed variation, including; diet, lifestyle, and health condition. Another major influence on the microbiome is the host’s genetic background. We hypothesized the immune system to be one of the most important genetic factors driving the differences observed between gut microbiomes. Therefore, we are interested in linking the polymorphic molecules that play a role in human immune responses to the composition of the microbiome. HLA molecules are the most polymorphic molecules in our genome and therefore makes an excellent candidate to test such an association/link. To our knowledge for the first time, our results indicate a significant impact of the HLA on the human gut microbiome composition.

## Introduction

The gut microbiota is essential for the existence of the mammalian host and performs crucial functions, such as nutrition and colonization resistance against pathogens (Kamada, 2013; Lawley & Walker, 2013). In addition, commensal bacteria shape host immunity (Hill & Artis, 2010; Renz et al., 2012): Commensal bacteria influence the development and homeostasis of the immune system, and in the absence of a gut microbiome the innate and adaptive immune system is impaired (Sonnenberg & Artis, 2012). These functions are fulfilled by the 10^13^–10^14^ bacteria making-up the human gut microbiome of a typical adult (Sender et al., 2016). The composition of the gut microbiota varies substantially over the human population (The Human Microbiome Project Consortium, 2012; Eckburg et al., 2005), and numerous exogenous and intrinsic factors are proposed to influence the microbial community composition leading to observed microbiome diversity (Falony et al., 2016; Zhernakova et al., 2016). These factors include dietary components, mode of delivery during birth, breastfeeding, disease history, and host genetics (Belkaid & Hand, 2014; Bonder et al., 2016; Goodrich et al., 2016, 2014; Kolde et al., 2018). Studies with monozygotic twins have highlighted the importance of host genetics in shaping the microbiome (Reyes et al., 2010; Zhou et al., 2016), where the immune system is thought to be one of the most important contributing factors (Khan et al., 2019).

In recent years, numerous factors have been shown to cover a part of the interindividual variation observed in the microbiome (Zhernakova et al., 2016). However, the largest section of population diversity is still unexplained. To obtain a better understanding of the microbiome composition and functionality, a comprehensive list of associated factors is a requirement. A well-known but not fully explored mechanism influencing the microbiome is the immune system (Khan et al., 2019; Mao et al., 2018; van Baarlen et al., 2009). Conversely, the microbiota also imprints its signature on the host immune system, e.g. by influencing the abundances of mucosal-associated invariant T cells by exposure to microbial antigens (Constantinides et al., 2019). Immune response related factors that generate variation between microbiomes are likely to come from polymorphic immune molecules, as they also vary between individuals.

Major Histocompatibility Complex (MHC) genes, known as the human leukocyte antigen (HLA) genes in humans, are one of the most polymorphic genes found in vertebrates (Mungall et al., 2003; Robinson et al., 2015). They encode cell surface glycoproteins that present antigens to T cells: the class I molecules present intracellular antigen to CD8+ T cells (Rock et al., 2016), while the class II HLA molecules present extracellular antigen to CD4+ T cells. Through cross presentation, the display of extracellular antigen on MHC class I is also observed (Heath & Carbone, 2001; Joffre et al., 2012). Together these molecules form the first essential step in the generation of an adaptive immune response. Influence of the MHC haplotype on the microbiome has been observed in mice (Khan et al., 2019; Kubinak et al., 2015; Lin et al., 2014; Toivanen et al., 2001) and in stickleback (Bolnick et al., 2014). In humans, specific polymorphisms in the HLA region have been associated to several common infections (Tian et al., 2017) and in a recent genome-wide association study of over 1,500 individuals two out of the nine genetic variants associated to the microbiome composition were within the HLA region (Bonder et al., 2016). An association with microbiome composition has been observed for the HLA-DQ gene in relation to Coeliac disease (Palma et al., 2010; Olivares et al., 2015).

Here we analyse 3,002 human gut microbiome datasets and find that (i) HLA zygosity correlates with microbiome diversity, (ii) HLA haplotype similarity correlates with microbiome diversity. Importantly, functionally stratifying individuals on the basis of HLA-presented peptides improved the association with microbiome similarity. Overall, these results provide the first statistical support for the functional association between HLA and the gut microbiome in humans.

## Results and discussion

### Mining gut microbiome datasets

To link the HLA haplotype to microbiome composition of individuals, we simultaneously determined the presence of human HLA genes and bacterial small subunit (SSU) rRNA genes in human intestinal samples. We composed a combined reference database consisting of HLA gene sequences and SSU ribosomal RNA sequences and used this to map 3,002 gut microbiome datasets extracted from the sequence read archive (SRA). As expected, we observed a large diversity in the number of HLA reads recovered from the different datasets (Supplementary Figure 1, Supplementary Table 1 and 2). This could be due to differences in sequencing techniques between studies or individual variation. For 127 and 309 individuals, we could predict the complete HLA haplotype for the classical HLA class I (HLA-A, HLA-B and HLA-C) and HLA class II (HLA-DP, HLA-DQ and HLA-DR) genes, respectively (Figure 1, details of this procedure are explained in the Methods section).

**Figure 1.**
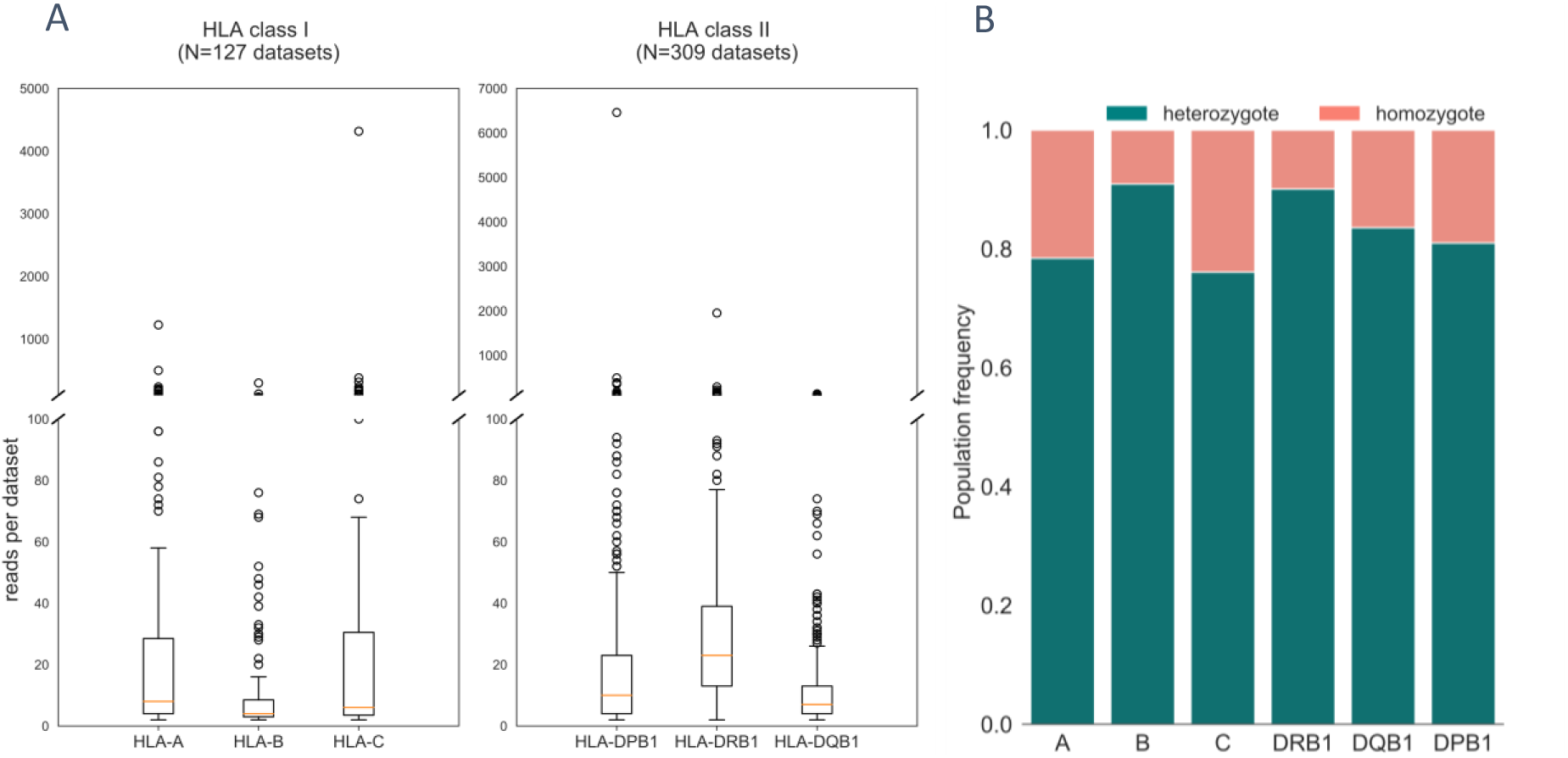
Overview of the dataset content per group selected on having a complete profile for the HLA class I or II genes. (A, left) Discontinuous bar plot showing the number of reads mapping to HLA-A, B or C genes for the datasets in the HLA class I group. (A, right) Reads mapping to HLA-DPB1, DRB1 or DQB1 genes for the datasets in the HLA class II group. (B) Relative frequency of homozygotes (pink) or heterozygotes (blue) for A, B, C, DRB1, DQB1 and DPB1 HLA genes.

Within the successfully HLA-haplotyped groups, most HLA reads were mapped to the HLA-A, HLA-C and HLA-DRB1 genes (Figure 1A). HLA class I is expressed in all nucleated cells; while HLA class II expression is restricted especially to antigen presenting cells (APCs) (Rock et al., 2016). Therefore, we expected a higher number of reads mapping to HLA class I in human gut datasets, but the difference between HLA class I and II reads is not significant (P=0.17, two-sample KS test). The observation of similar read counts between HLA class I and II may be explained by a large number of lymphocytes being present in the gut as a part of mucosal immunity (McDermott & Huffnagle, 2014). Moreover, intestinal epithelial cells also express MHC class II (Wosen et al., 2018), and HLA class I and class II expression levels vary across different cell types (Handunnetthi et al., 2010; Johnson, 2000). Further differences in the gene expression pattern of gut associated cell types, as well as stochastic differences in e.g., sampling and analysis protocols, might contribute to the observed variation in HLA mapped reads between the SRA datasets. We observed a high frequency of HLA heterozygosity and low frequency of homozygosity for any of the HLA genes (Figure 1B), consistent with the high polymorphism of HLA genes in the population (Maiers et al., 2007).

### Inferred HLA prevalence reflect the Caucasian population profile

As a quality assessment of our HLA typing method from gut microbiome datasets, the population allele frequencies of the HLA class I and class II groups were compared to the HLA prevalence in the National Marrow Donor Program (NMDP) database, a database containing high-resolution HLA allele frequencies (Maiers et al., 2007). The frequency distribution of HLA class I in our data set is similar to that in the NMDP database (Pearson’s r=0.806, r=0.726, r=0.812, HLA-A, HLA-B, and HLA-C, respectively), but the frequency distribution of HLA class II less so (Pearson’s r=0.537, r=0.127, HLA-DQB1 and HLA-DRB1, respectively) (Supplementary Figure 4A-E). All genes except HLA-C had zygosity frequencies that were consistent with the NMDP data (Supplementary Figure 4F). Together, the HLA haplotypes (especially for class I molecules) predicted from public human gut microbiome samples showed a similar allele frequency distribution and zygosity when compared to the NMDP data, providing support for our HLA typing approach.

### Microbiome alpha diversity is higher in individuals with homozygous HLA alleles

To investigate how HLA diversity and microbiome diversity are related, we tested first the effect of HLA heterozygosity on the alpha diversity of the gut microbiota. For this analysis, we used a subset of datasets of which we could reliably determine the full zygosity of all HLA class I or II genes (n=60 and n=205, respectively). Although we found only very few individuals who were fully homozygous for all HLA genes, we observed a significantly higher microbiome richness in homozygous individuals than in individuals with one or more heterozygous HLA class I genes (Figure 2A). Moreover, we observed a trend for a decrease of the richness as the homozygosity decreases (Figure 2A). Similar results were obtained for HLA class II (Supplementary Figures 5E-F). We also investigated the correlation between HLA zygosity and microbiome evenness, but this yielded inconsistent results (Supplementary Figure 5A-D).

**Figure 2.**
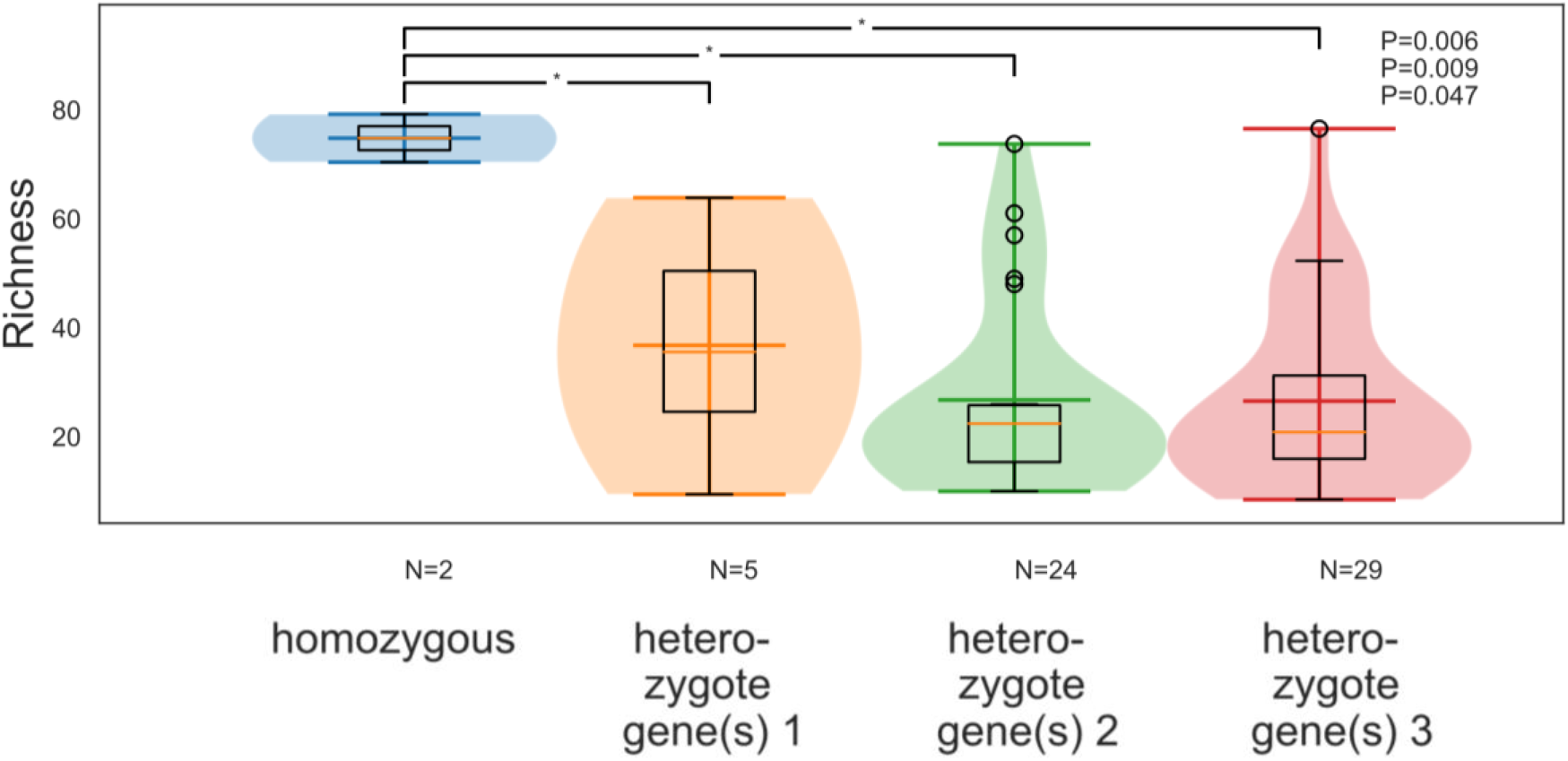
The influence of HLA zygosity on gut microbiome alpha diversity. Violin box plots of the Margalef’s richness index for datasets grouped on the number of heterozygote HLA class I genes. Hypothesis one-sided two sample Kolmogorov-Smirnov test: individuals with 0 heterozygote genes richness is higher than other groups. P-values of Kolmogorov-Smirnov test between the other groups in Supplementary Figure 5G.

Next, we analysed whether the zygosity effect on alpha diversity is equally distributed over the HLA class I genes. For HLA class I genes, individuals homozygous for HLA-B genes had a higher richness compared to HLA-A and HLA-C (Figure 3). We did not find a significant difference among the HLA class II genes (Supplementary Figure 6A). Association between the gut microbiome and HLA-B was observed previously (Bonder et al., 2016; Lin et al., 2014) and is consistent with HLA-B being the immunodominant MHC class I gene (Bihl et al., 2006; Lacey et al., 2003; Lewinsohn et al., 2007; Weiskopf et al., 2011) and the most polymorphic one (Robinson et al., 2015). We also observed a higher microbiome evenness associated with HLA-DQB1 homozygosity (significant) and HLA-B homozygosity (trend) (Supplementary Figure 6B-C).

**Figure 3.**
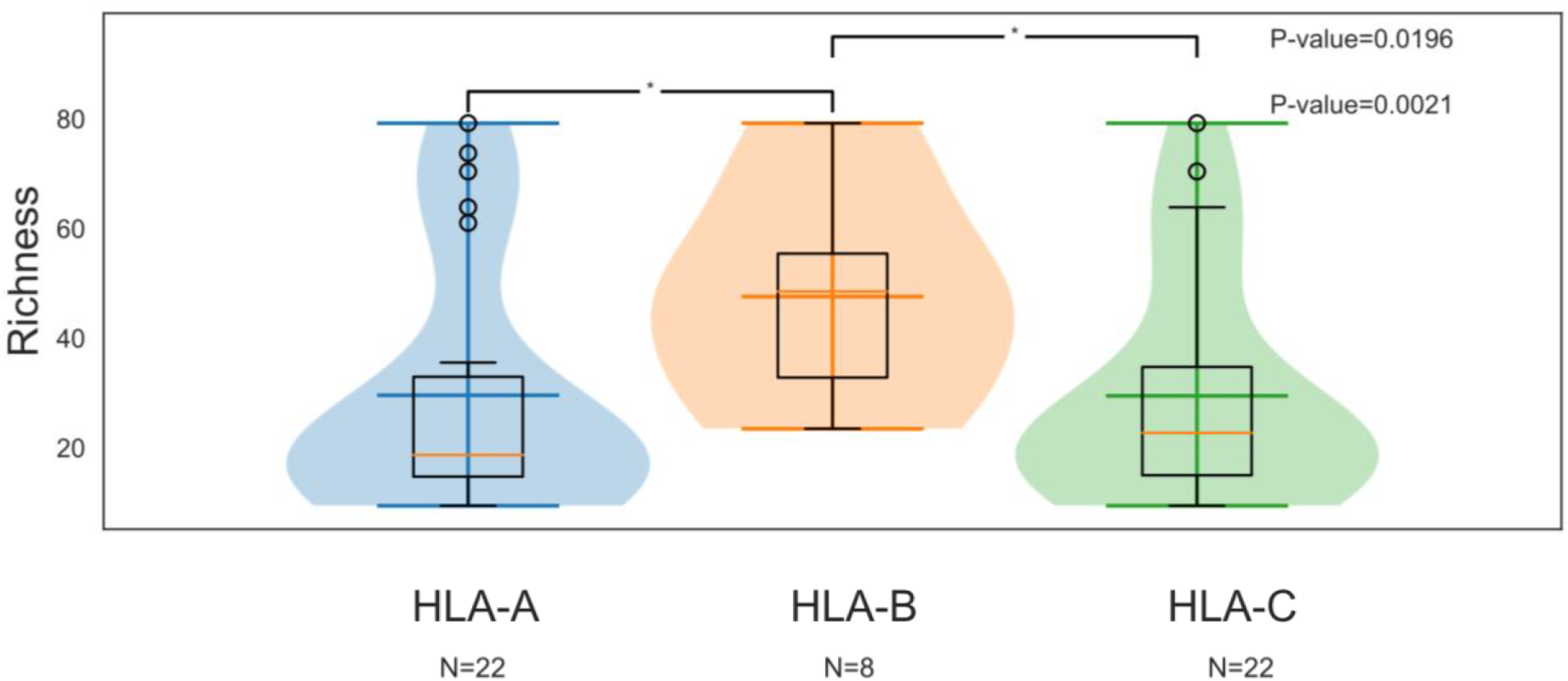
The influence of homozygosity of different HLA class I genes on the gut microbiome alpha diversity. Richness for individuals who are homozygote for HLA-A (blue), HLA-B (orange), or HLA-C (green) class I genes. HLA-B richness is higher than HLA-A or HLA-C (two-sided two sample KS-test); HLA-A and HLA-C do not differ significantly in Richness (two-sided two sample KS-test, P-value= 0.8717).

### Individuals with similar HLA genes have similar microbiomes

We next asked if a higher microbiome similarity was observed in individuals who shared more HLA alleles. To do this, we determined the similarity in microbiome composition for pairs of datasets that were stratified by the number of shared HLA class I alleles (Supplementary Figure 7). As expected, microbiome similarity was generally higher between individuals who shared more HLA alleles, although the effect size was modest, possibly as a result of the low number of sample pairs with highly similar HLA profiles.

We expect that two individuals with the same HLA molecules may have similar immune control of their microbiome. However, the opposite is not true, i.e., individuals with different sets of HLA molecules can nevertheless generate similar immune responses because different HLA molecules may have similar binding motifs. To address this issue of functional similarity, HLA molecules have been grouped into supertypes, consisting of HLA molecules that present similar peptides on the cell surface (Sidney et al., 2008). In a first attempt to represent the functional similarity in HLA between individuals and increase statistical power at this highly polymorphic gene, we performed association analyses at the level of HLA supertypes. While this increased the number of sample pairs with similar HLA profiles, the relation between HLA zygosity and microbiome alpha diversity was lost (Supplementary Figure 8). This may be because the HLA supertypes are a very generalized representation of HLA molecules, while the functional effects of the HLA molecules on the microbiome composition are likely very specific (van Deutekom & Keşmir, 2015).

In a second attempt to quantify the functional similarity with respect to HLA between individuals, we devised a novel approach by focusing on the peptides presented by the HLA molecules. To do this, we used NetMHCpan4.0, a state-of-the-art peptide-MHC binding affinity prediction tool (Jurtz et al., 2017) to predict the binding affinity of over two million gut bacterial oligopeptides to the HLA class I molecules. The peptides were extracted from the human gut catalog (Li et al., 2014) in order to best represent the HLA functionality in the context of their potential response to the human gut microbiome. This approach allowed us to calculate a Presented Peptidome Similarity Score (PPSS) between the HLA haplotypes of each pair of individuals (see Methods). Individuals with a high PPSS present very similar peptides, while individuals with a low PPSS present different peptides. When we stratified pairs of individuals by PPSS, we observed a significantly higher microbiome similarity for pairs with high PPSS (Figure 4).

**Figure 4.**
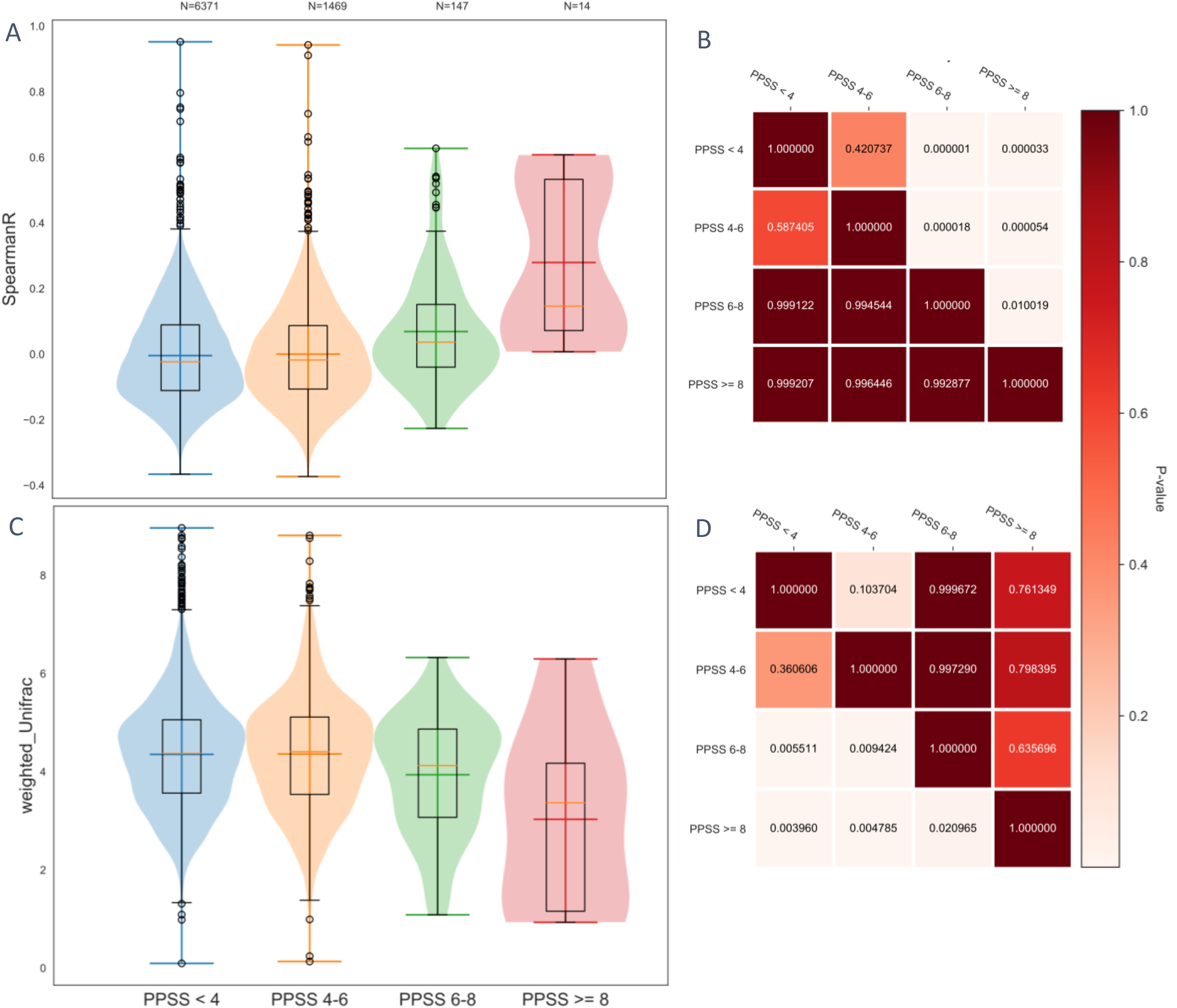
Individuals presenting similar microbial human gut peptides on their HLA have a similar microbiome. Spearman Rank correlation (A) and weighted UniFrac distance (C) of the microbiome composition for sample pairs with <4 PPSS (Blue), 4-6 PPSS (orange), 6-8 PPSS (green) and PPSS >= 8 (red) scores. (B and D) Heatmap displaying the P-values for the two sample Kolmogorov-Smirnov test under the null hypothesis; item on the y-axis is drawn from an equal or smaller distribution than the item on the x-axis.

The mechanism by which HLA class I molecules might influence the gut bacterial composition remains unclear. Little is known about how MHC molecules affect the gut microbiome. MHC class II molecules are suggested to play a role via antibody-mediated response (Bolnick et al., 2014; Khan et al., 2019; Kubinak et al., 2015b). One possibility is that a less diverse HLA repertoire in homozygous individuals may reduce the number and diversity of immune responses to the microbiota by T cells and natural killer (NK) cells. As a result, homozygous individuals would recognize and potentially remove fewer bacterial species from their intestinal microbiome than heterozygous individuals. This could allow a higher number of different bacterial species to colonize the gut than the individuals with a more diverse set of HLA molecules. A similar effect has been observed in the stickleback, where fish with more diverse MHC class II molecules had a less diverse gut microbiome (Bolnick et al., 2014). We focused our analysis on HLA class I molecules because the imputed HLA class II haplotypes did not match the expected population profile (Supplementary Figure 4), but we expect to find similar results with HLA class II haplotypes.

HLA class I molecules are involved in intracellular antigen presentation, leading to a cytotoxic response by CD8+ T cells. Cross-presentation of extracellular antigens on MHC class I by dendritic cells is very common (Heath & Carbone, 2001; Houde et al., 2003; Joffre et al., 2012; Kamphorst et al., 2010), but it remains unknown how antigen presented on HLA class I shapes mucosal immunity. In addition, the influence of HLA class I on the microbiome could be related to the MHC class I chain-related antigen A (MIC A) and antigen B (MIC B) genes. These polymorphic genes are constitutively expressed in the gut epithelium (Groh et al., 1996). The recognition of MIC-A/B by its ligand NKG2D, an activating receptor on NK cells, NKT cells, and T cells, leads to cytotoxic response and cytokine production (Bauer et al., 1999). MIC-A/B are localized next to HLA-B in chromosome 6, and therefore in strong linkage disequilibrium with HLA-B (Ando et al., 1997; Petersdorf et al., 1999). Because we found the strongest homozygosity effect on the gut microbiome for HLA-B (Figure 3), it will be of interest to dissect the association between HLA-B or MIC-A/B and the microbiome in large, well-controlled cohorts (Falony et al., 2016; Wijmenga & Zhernakova, 2018; Zhernakova et al., 2016).

Above, we presented the first statistical evidence in humans that HLA significantly affects the composition of the gut microbiota, in line with previous results in other animals (Khan et al., 2019; Kubinak et al., 2015; Mao et al., 2018). We expect that the immune system may be one of the most important genetic factors influencing the microbiome (Bonder et al., 2016; Goodrich et al., 2014). As HLA is a readily measurable parameter of the immune system, the exploration of HLA and HLA-driven peptide presentation opens the door for incorporating these factors into future predictive models of the gut microbiome (Falony et al., 2016; Zhernakova et al., 2016).

## Materials and Methods

### Human gut microbiome datasets

We retrieved 3,002 human gut datasets from 37 different Bio Projects from the public Sequence Read Archive (SRA) database (Leinonen, Sugawara, Shumway, & International Nucleotide Sequence Database Collaboration, 2011). Datasets were collected using “metatranscriptome” search terms (Table 1), but due to inconsistencies in SRA metadata annotation, our selection included 750 metagenomes as well. Microbiome and HLA haplotype profiling are performed on both data types. Repeating the analyses with the subset of 2,250 metatranscriptomes, led to a steep decrease in the number of paired individuals with highly similar HLA profiles and loss of signal, and therefore we decided to continue with the original dataset. Metadata fields are listed in Supplementary Tables 1-2. A full list of dataset identifiers may be requested from the authors. We are grateful to the associated studies for making these data available for reuse.

**Table 1.** Search terms SRA query. The search terms used are listed in column 1, columns 2-4 are the selected filters, and column 5 is the number of returned datasets in the SRA database. Overlapping datasets from the different search terms were filtered out.

| Search term | Filter [Source] | Filter [Organism] | Filter [Selection] | Number of datasets |
| --- | --- | --- | --- | --- |
| Human bowel | Metatranscriptomic | - | Random | 841 |
| Human gut | Metatranscriptomic | - | Random | 925 |
| - | Transcriptomic | human gut metagenome<br>orgn txid:408170 | Random | 130 |
| - | Metatranscriptomic | human gut metagenome<br>orgn txid:408170 | Random | 897 |
| Human stool | Metatranscriptomic | - | Random | 778 |
| RNA sequencing | - | human gut metagenome<br>orgn txid:408170 | Random | 2,879 |
| Metatranscriptomic | - | human gut metagenome<br>orgn txid:408170 | - | 2,179 |

All sequence runs were pre-processed as follows. Datasets were downloaded using prefetch (version 2.9.2) utility and SRA data was converted to fastq files using fastq-dump (version 2.9.2), both from the SRA toolbox. The quality of the reads was assessed using FastQC (version 0.11.8, S. Andrews, 2010) and reads where trimmed using AdapterRemoval (version 2.1.7, Schubert et al., 2016). For studies with multiple datasets from the same individual, we randomly selected one dataset to include in our analysis to avoid biasing the dataset with intra-individual sample comparisons.

### Mapping reads to a composite reference database

To assess the microbiome composition and the HLA haplotype at once, reads were mapped with BWA-MEM (version 0.7.12, H. Li & Durbin, 2009) to a reference database consisting of bacterial small subunit (SSU) rRNA (SILVA Release 132, Ref NR 99, Quast et al., 2012) and genomic HLA sequences from IMGT/HLA database (Release 3.31.0, Robinson, et al., 2015). 16S reads were analysed to obtain the bacterial composition of each dataset. Only bacterial OTUs with a relative abundance ≥0.0001 were taken into account. Successively, the datasets should contain ≥1,000 16S reads. We developed a sorting scheme for multi-mapping reads, which were divided across target sequences as a fraction of the number of primary mapping (unique) reads (Clausen et al, 2018; Coutinho et al., 2017; Iverson et al., 2012).

### HLA imputation

We performed two-digit HLA haplotyping on the classical HLA loci (HLA-A, HLA-B, HLA-C, HLA-DR, HLA-DP and HLA-DQ). Although we expect a maximum of two alleles for each gene (Rock et al., 2016), we often found more than two HLA alleles with reads mapped. This is likely a result of the high sequence similarity of the different alleles from the same HLA gene. In order to make most reliable HLA imputation we defined an R score as the ratio of the number of reads mapped to the two most frequently observed HLA alleles over the number of reads mapped to all other alleles of this gene (Figure 5A). We determined R for each of the six classical HLA loci in all 3,002 datasets. Only HLA genes with an R score of two or higher were considered for HLA imputation (see Figure 5B). An individual was considered homozygotic at a given locus if at least ten reads aligned to the top HLA allele and the second mapped allele had less than one fifth of the reads of the first. A locus was considered heterozygotic if at least two alleles were found and the second mapped at least one fifth of the reads of the first.

**Figure 5.**
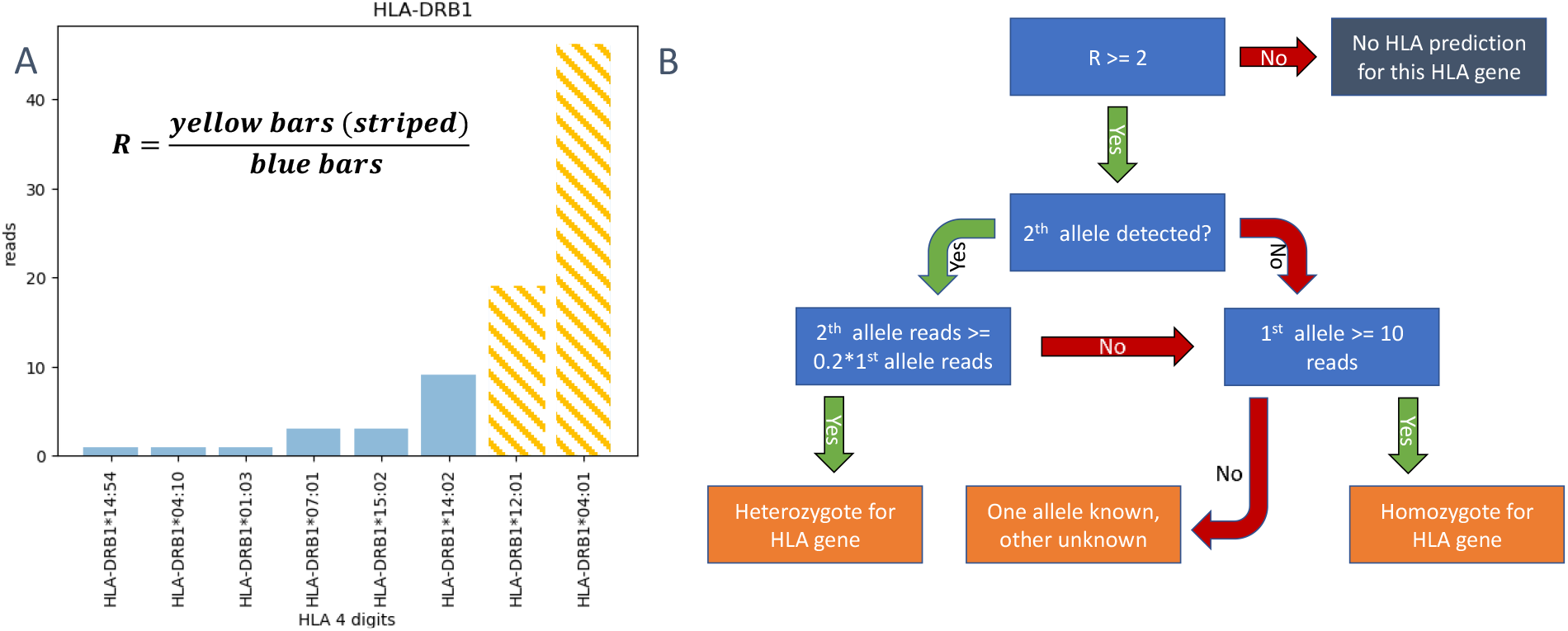
Decision tree genotyping HLA gene. (A) Visual explanation of the R score. (B) Decision tree for HLA haplotyping from fecal microbiome datasets. Blue boxes are the decisions, orange boxes represent possible positive outcomes and the grey box is the negative outcome when no prediction is made for this gene.

### Dataset selection and population-based HLA haplotyping quality control

Individual HLA loci may be in linkage disequilibrium with other loci (Hiller et al., 1978; Jeffreys et al., 2001). To maximize the representation of the HLA profile, we only included datasets if the complete HLA haplotype was assessed, resulting in two groups for the classical class I and II (n=127, n=309, respectively, see Supplementary Tables 1 and 2).

Quality control of our haplotyping approach was performed by comparing the population allele and zygosity frequencies of our data with the NMDP ‘High-Resolution HLA Alleles and Haplotypes in the US Population’ database (Maiers et al., 2007). Data is extracted in September 2019. Based on the dataset localities, we expect a majority of individuals in our datasets to be from European descent (see Supplementary Table 1 and 2). Therefore, we compared our HLA frequencies with the cohort of European origin in NMDP database.

### HLA supertypes and Presented Peptidome Similarity Score (PPSS)

Most of the 450 haplotyped datasets had unique HLA profiles, reflective of the high population-level diversity at the HLA loci. To increase statistical sensitivity, we performed two separate experiments on the data. First, we clustered HLA-A and HLA-B alleles into six supertypes each (Sidney et al., 2008). Second, we devised a Presented Peptidome Similarity Score (PPSS) between two haplotypes to estimate the functional similarity in immune profile, based on the overlap in peptides presented by their HLAs. To do this, first we created a peptidome dataset consisting of human gut bacterial peptides sampled from the 9,878,647 proteins in the human gut catalogue (Li et al., 2014). By using a sliding window of length nine amino acids, we randomly included one in every thousand oligopeptides into the peptidome and removed duplicates, resulting in a peptidome of 2,371,920 peptides. Next, we used NetMHCpan4.0 to predict which peptides were presented by each HLA class I allele (Jurtz et al., 2017). We used the recommended cut-off of 2.0 %Rank for classifying binding peptides.

The PPSS between two individuals *i* and *j* is defined as the summed Jaccard indices *J_g,ai,aj_* of the sets of peptides presented by all combinations of the alleles *ai* and *aj* of HLA gene *g* (HLA-A, HLA-B, and HLA-C). PPSS ranges from zero (no overlap between presented peptides) to twelve (presented peptides are identical). The Jaccard based similarity of peptide pools presented of allele pair a,b, is defined as the overlap of peptide pools (O_a,b_) divided by the not shared peptide pools (Eq. 1). The J_ab_ scores between all two-digit alleles are shown in Supplementary Figure 2. PPSS_ij_, the similarity of peptide pools of the pair of individuals i and j, is defined by the allele pair similarity for all combinations of alleles within one gene, added up for all three HLA genes (Eq.2). The distribution of PPSS scores across all datasets is shown in Supplementary Figure 3.

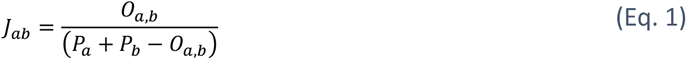

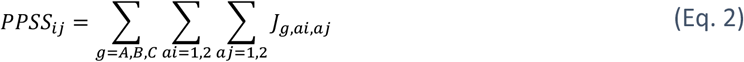

Supplementary Tables 3-5 contain the Jaccard indices for every pair of four-digit alleles for HLA gene HLA-A, HLA-B, and HLA-C, respectively, allowing readers to readily calculate the PPSS upon HLA haplotyping.

### Alpha diversity

To assess alpha diversity, we used evenness and richness measures from the skbio.diversity.alpha package (version 0.5.5, Scikit-bio development team, 2014). We did not rarefy the data, as this may increase type I and II errors (McMurdie & Holmes, 2014). We did not find a correlation between dataset size and microbiome richness (r=0.027, P=0.16, Pearson), and therefore we assume that dataset size does not bias our results. To approximate the number of different species in a sample, we used Margalef’s richness index (D_mg_) which attenuates sampling size effects (Eq. 3, Magurran, 2004), where S is the number of different OTUs observed in the sample and N is the number of 16S reads mapped:

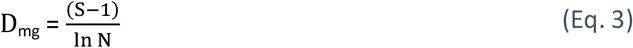

The microbiome evenness was calculated using Heip’s evenness measure (E) (Eq. 4, Heip, 1974), where H is the Shannon-Weiner entropy of the 16S reads and S is the number of OTUs in the dataset:

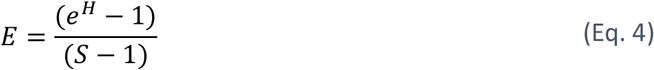

For the comparison between the alpha diversity and the number of heterozygote genes, only the fully classified datasets were used (Supplementary Table 1 and 2). In the analysis of the effect of homozygous HLA genes on alpha diversity, all datasets with at least one homozygous gene(s) were used.

### Beta diversity

The microbiome similarity between individuals was calculated using two beta diversity measures: weigthed UniFrac distance and Spearman’s Rank correlation (Lozupone et al., 2011). We adapted the SILVA taxonomy with a branch length of one between taxonomic ranks and used these distances for weighted UniFrac. The Spearman rank correlation was calculated using the abundances of the OTUs observed in each microbiome.

### Statistics

We used the Kolmogorov-Smirnov (KS) test for two samples from the python package scipy stats (version 1.3.2) for statistical analysis (Scipy Stats, 2019). With alternative hypothesis; ‘two-sided’, ‘less’ and ‘greater’ depending on the relevance for the comparison as stated in figure captions.

### Code Availability

Scripts for downloading the datasets, mapping the datasets to the composed reference genome, and further analysis can be accessed on github (https://github.com/Stijn-A/Quantifying_the_impact_of_Human_Leukocyte_Antigen_on_the_human_gut_microbiomes).

## Acknowledgements

We thank Prof. Dr. R.E. Bontrop for helpful discussions. B.E.D. was supported by the Netherlands Organization for Scientific Research (NWO) Vidi grant 864.14.004.

## Supplementary Figures

**Supplementary Figure 1.**
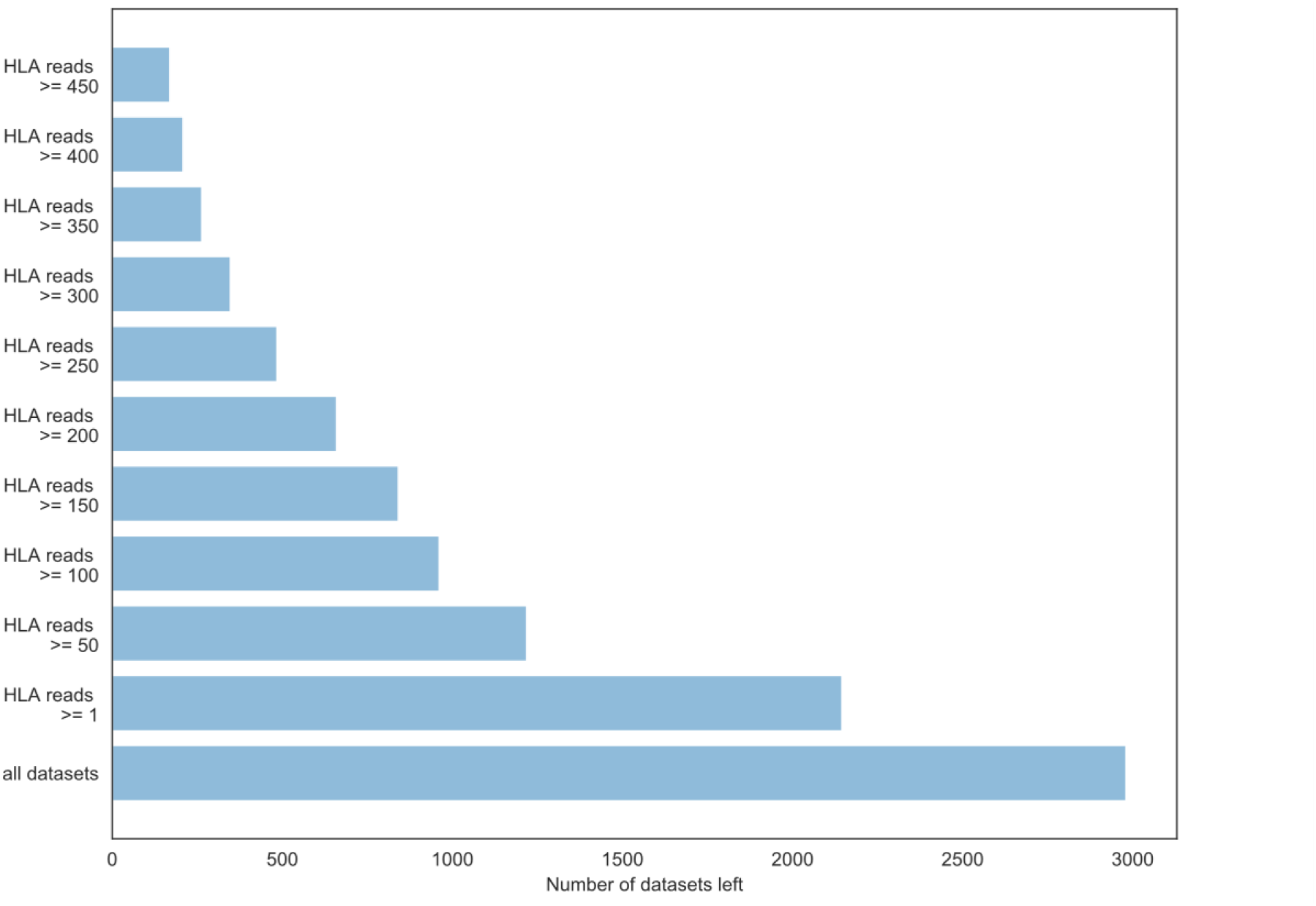
Reads mapping to HLA genes in all samples. The number of samples containing different numbers of HLA reads.

**Supplementary Figure 2.**
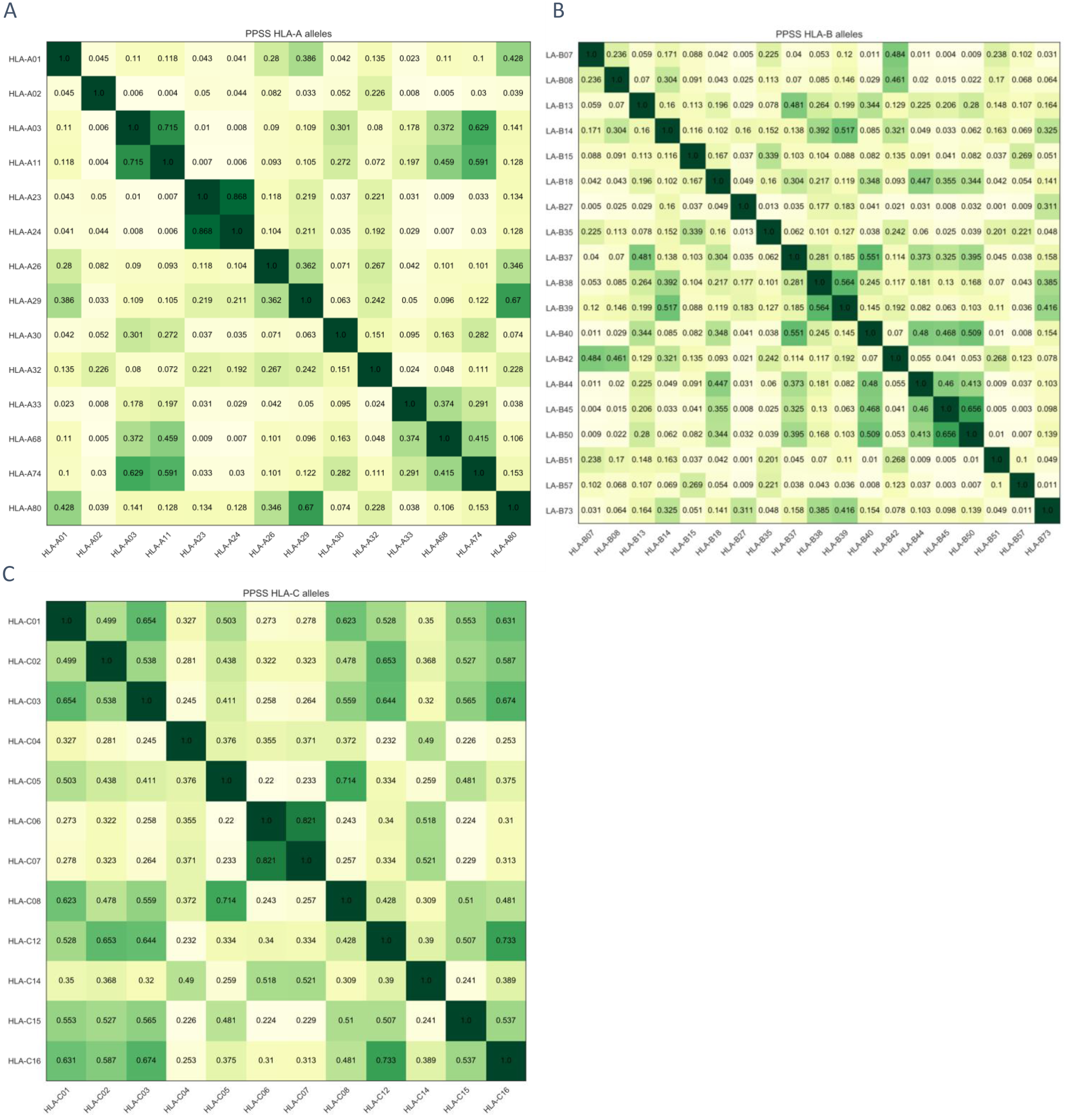
**Jaccard peptide presentation score between different HLA molecules** for HLA-A (A), HLA-B (B) and HLA-C (C) genes.

**Supplementary Figure 3.**
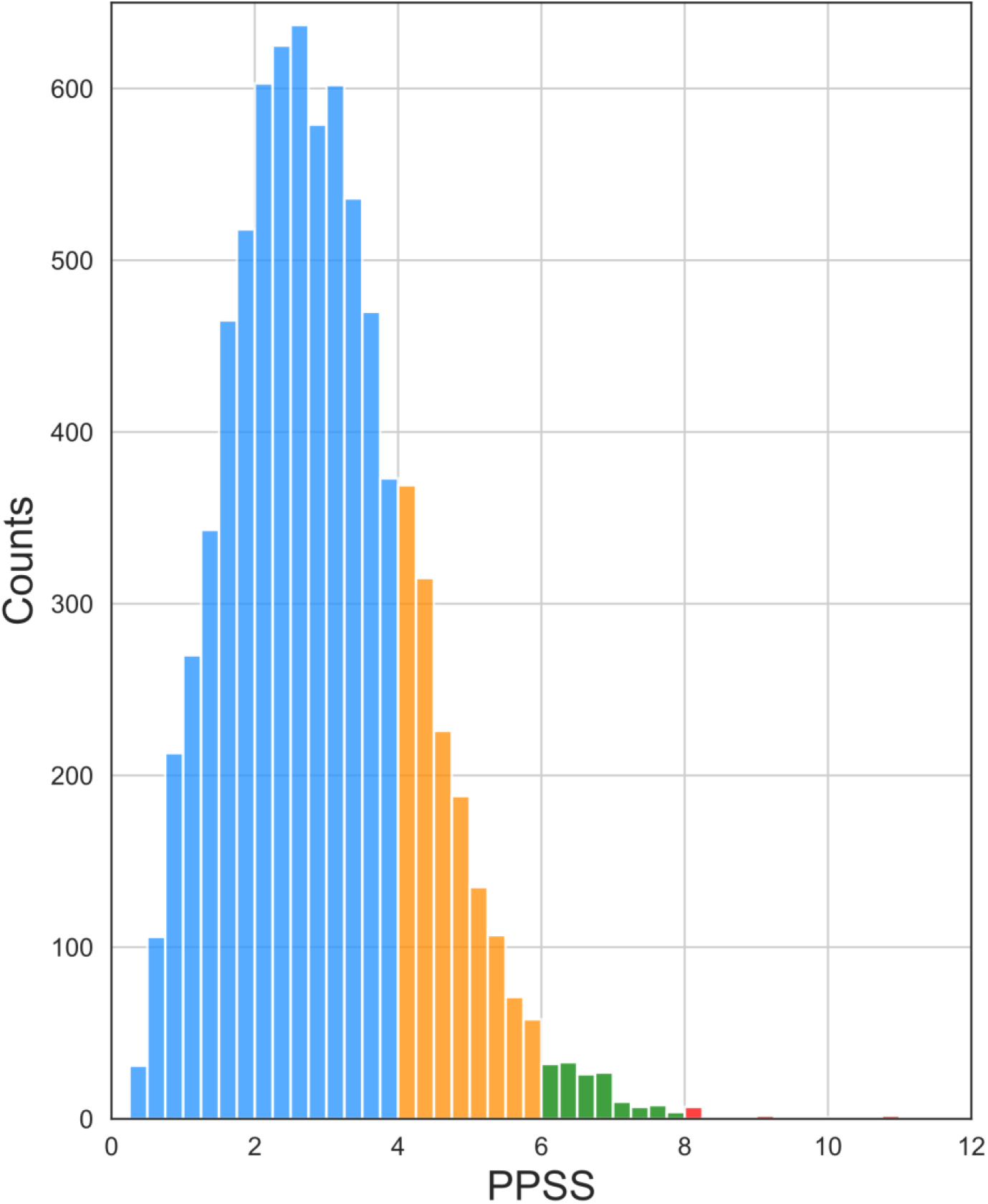
Histogram of PPSS pairs of individuals. The bar colors represent the stratification used in Figure 4; sample pairs with <4 PPSS (Blue), 4-6 PPSS (orange), 6-8 PPSS (green) and PPSS >= 8 (red) scores

**Supplementary Figure 4.**
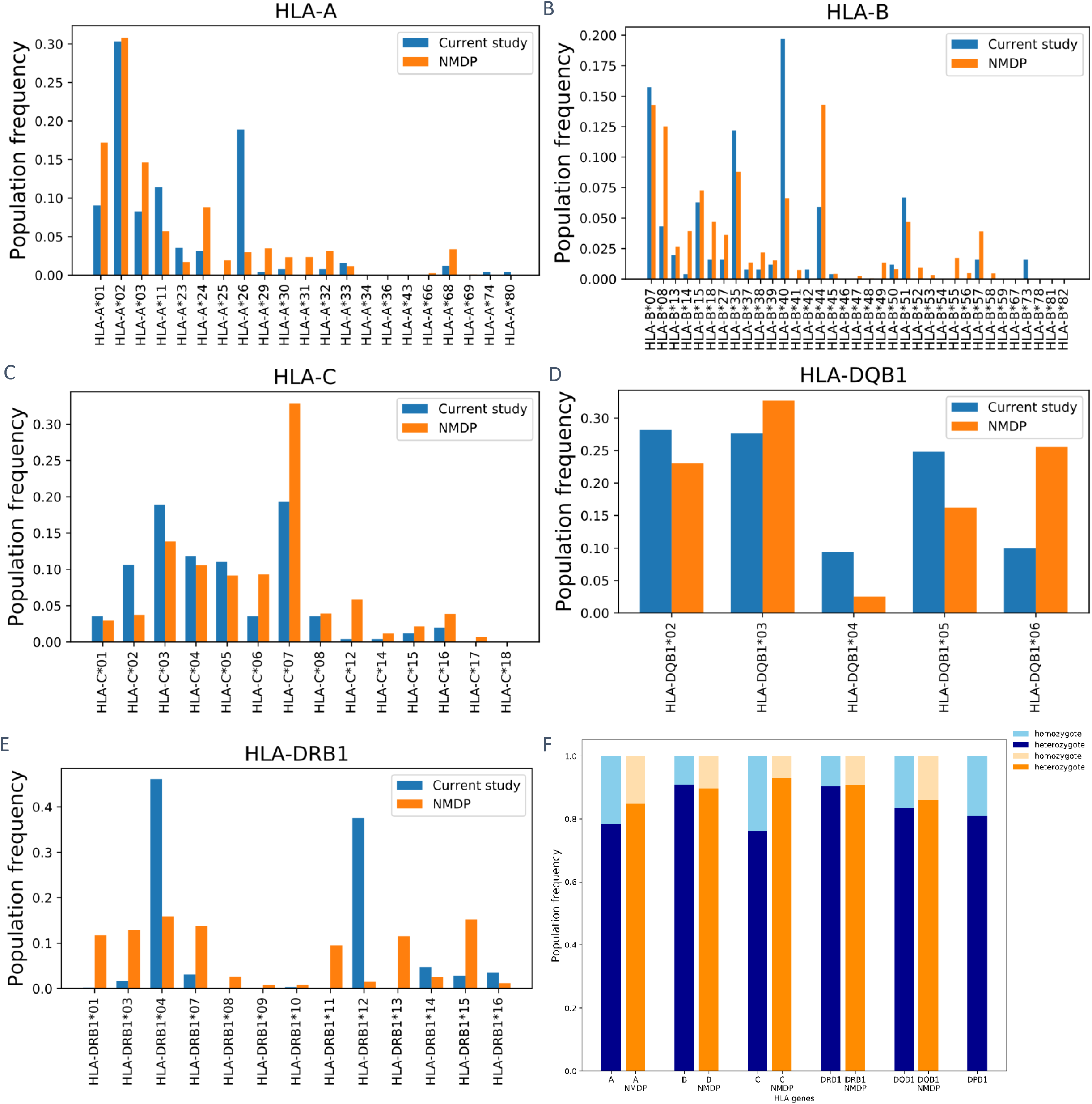
HLA frequencies compared between current study and National Marrow Donor Program (NMDP). (A to C) The HLA frequency distribution of the 127 samples selected on HLA class I compared to the HLA frequency of European origin in the NMDP database (Maiers et al., 2007), for: HLA-A (A), HLA-B (B) and HLA-C (C). Histogram of the distribution of the HLA class II DQB1 (D) and DRB1 (E) genes compared to the NMDP. The heterozygote/homozygote distribution for the HLA class I and II genes (F), again the data is compared to the NMDP zygosity frequencies. Data for the DPB1 gene was absent from the NMDP database.

**Supplementary Figure 5.**
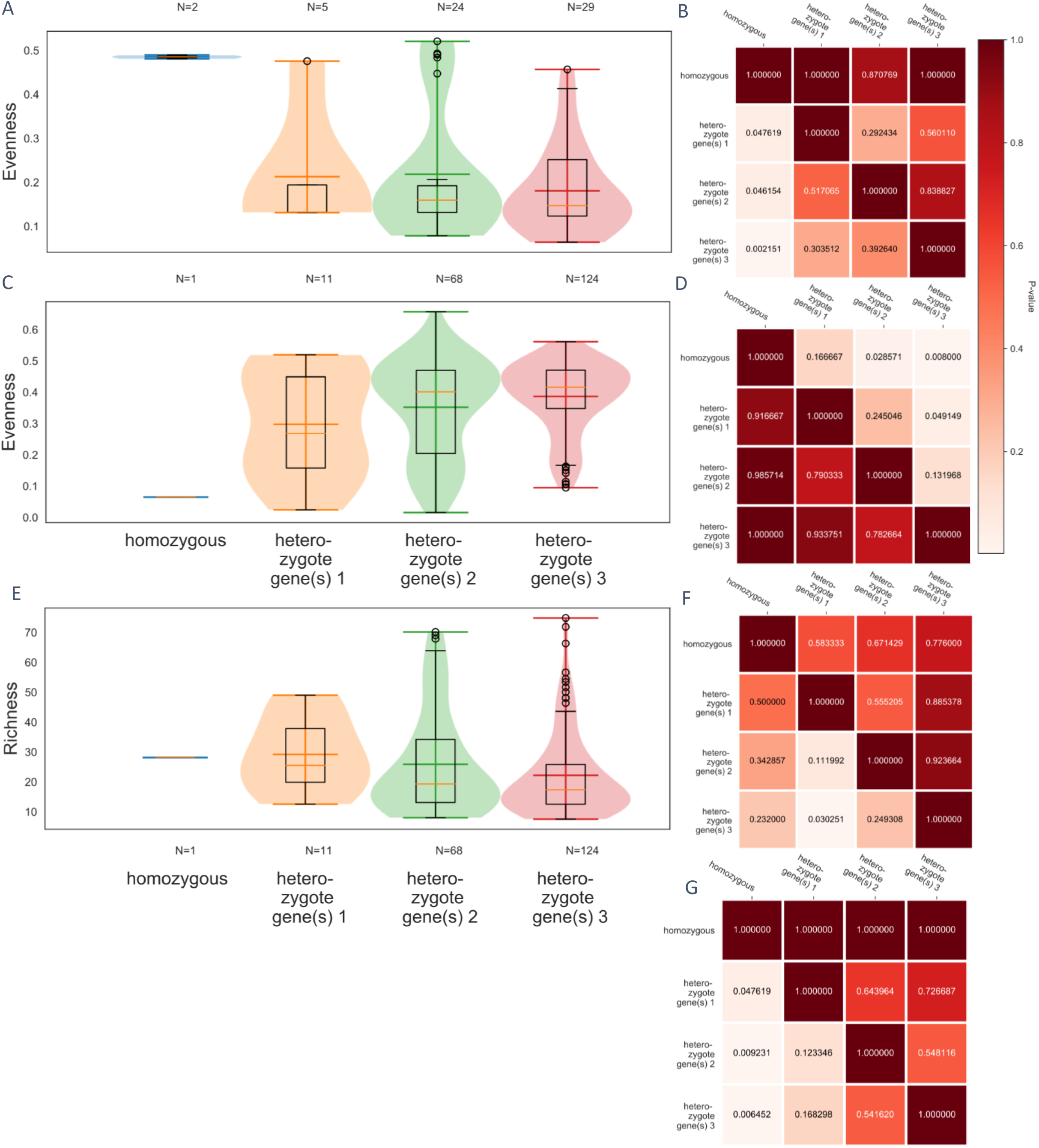
Alpha diversity over hetero- & homozygotic HLA class I and II genes. Violin box plots of the Heip’s evenness measure for individuals stratified on the number of HLA class I (A) and II (C) genes (Heip, C., 1974). (E) Violin box plots of theef’s richness index for datasets grouped on the number of heterozygote HLA class II (Magurran, 2004). (B, D, F, and G) Heatmaps on the right displays the P-values for the two Kolmogorov-Smirnov test under the null hypothesis; item on the y-axis is drawn from an or smaller distribution than the item on the x-axis. For the HLA class I evenness (C), HLA evenness (D), HLA class II richness (F) and the HLA class I richness (G).

**Supplementary Figure 6.**
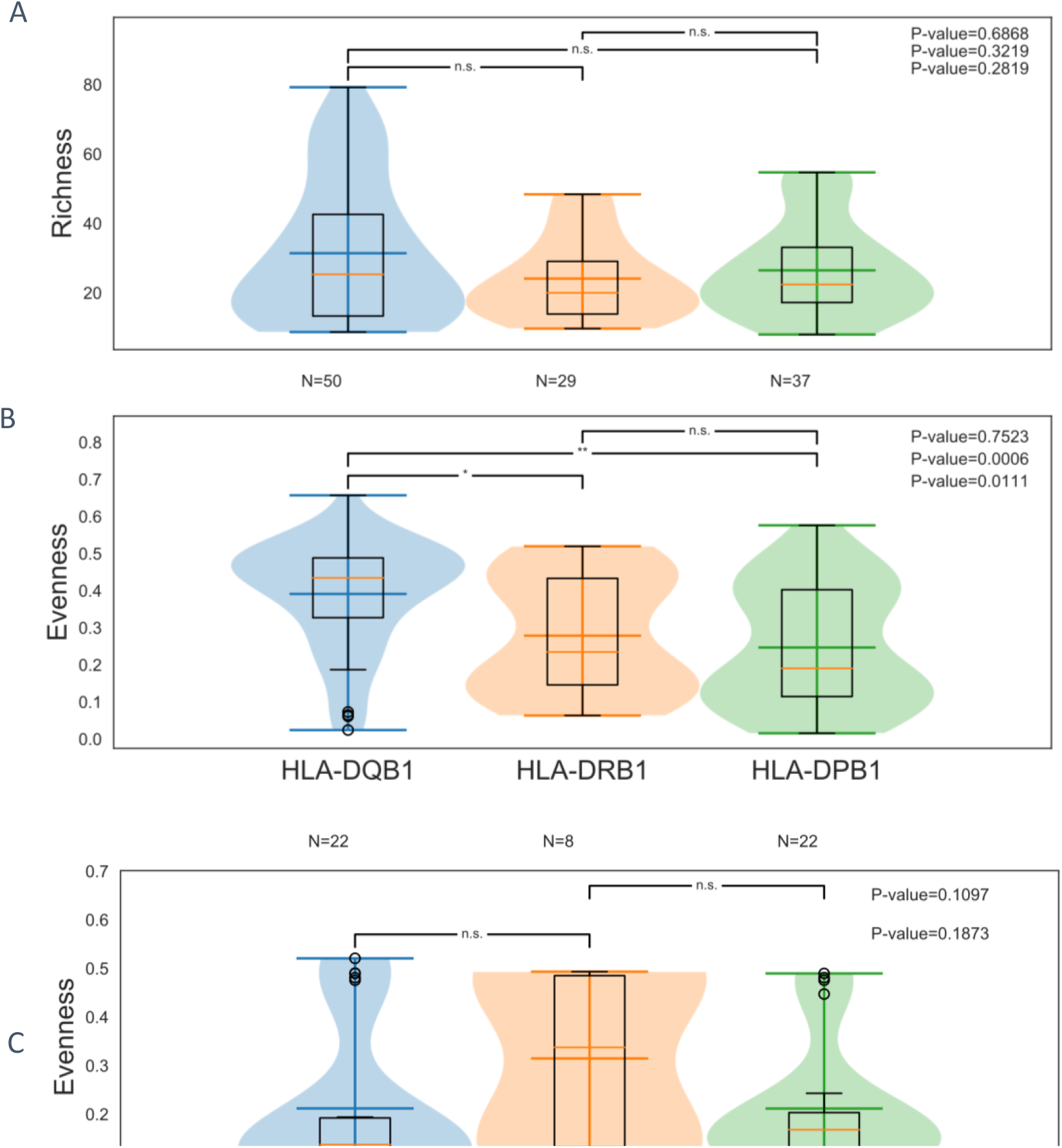
Effect of homozygous HLA genes on the alpha diversity of the microbiome. Richness (A) and Evenness (B) for individuals who are homozygote for HLA-DQB1 (blue), HLA-DRB1 (orange), or HLA-DPB1 (green) class II genes. P-values computed via two-sided two sample Kolmogorov-Smirnov test (n.s. P >= 0.05, * P < 0.05, ** P < 0.01, *** P < 0.001). (C) Evenness for individuals who are homozygote for HLA-A (blue), HLA-B (orange), or HLA-C (green) class I genes. HLA-A and HLA-C do not differ significantly in Richness or Evenness (two-sided two sample KS-test P-value= 0.5281).

**Supplementary Figure 7.**
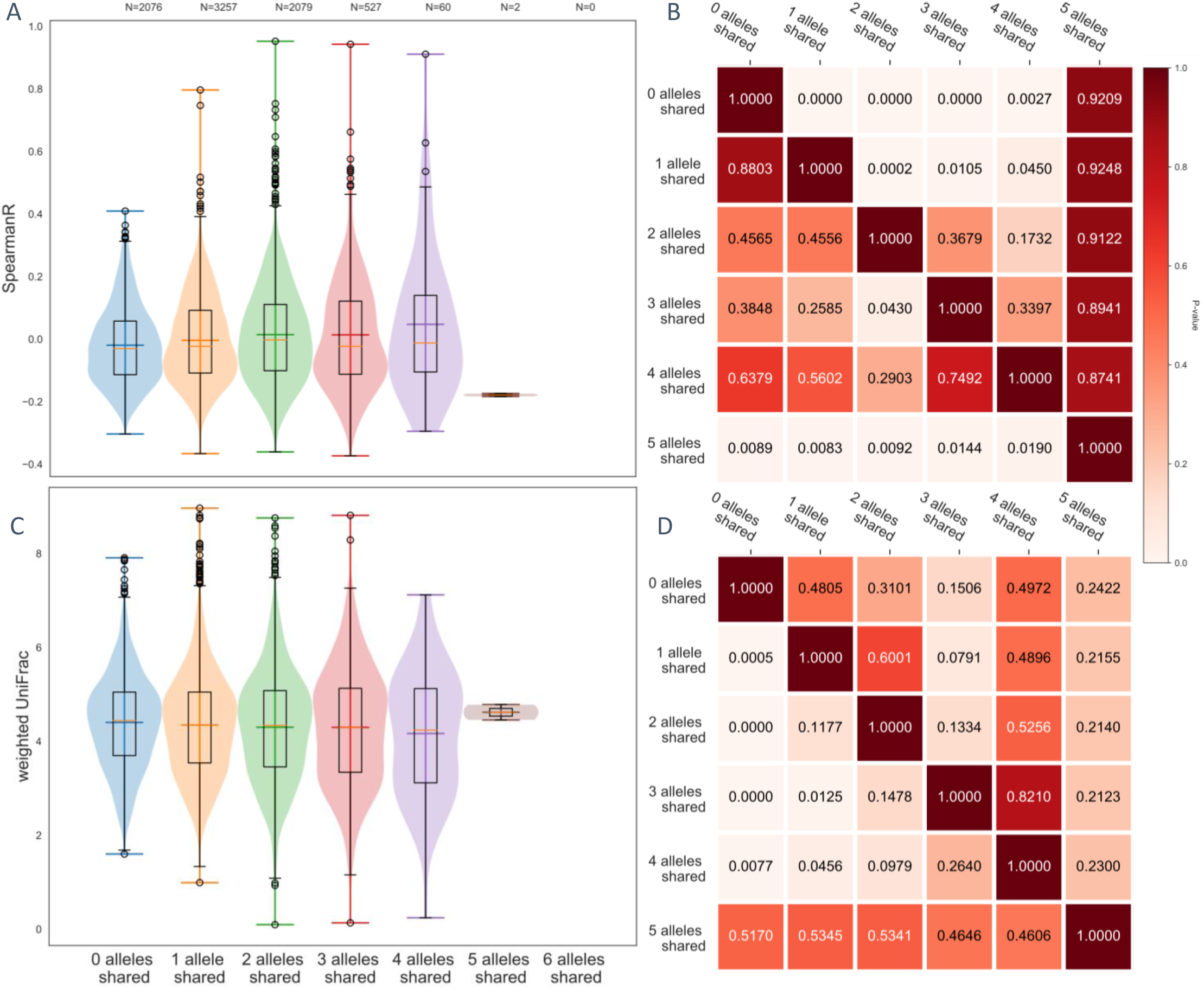
Microbiome beta diversity for samples with shared HLA class I alleles. (A) Spearman Rank correlation and (C) weighted UniFrac of the microbiome composition for sample pairs with 0 (blue), 1 (orange), 2 (green), 3 (red), 4 (purple), or 5 (brown) alleles shared. (B and D) Heatmaps on the right displays the P-values for the two sample Kolmogorov-Smirnov test under the null hypothesis; item on the y-axis is drawn from an equal or smaller distribution than the item on the x-axis.

**Supplementary Figure 8.**
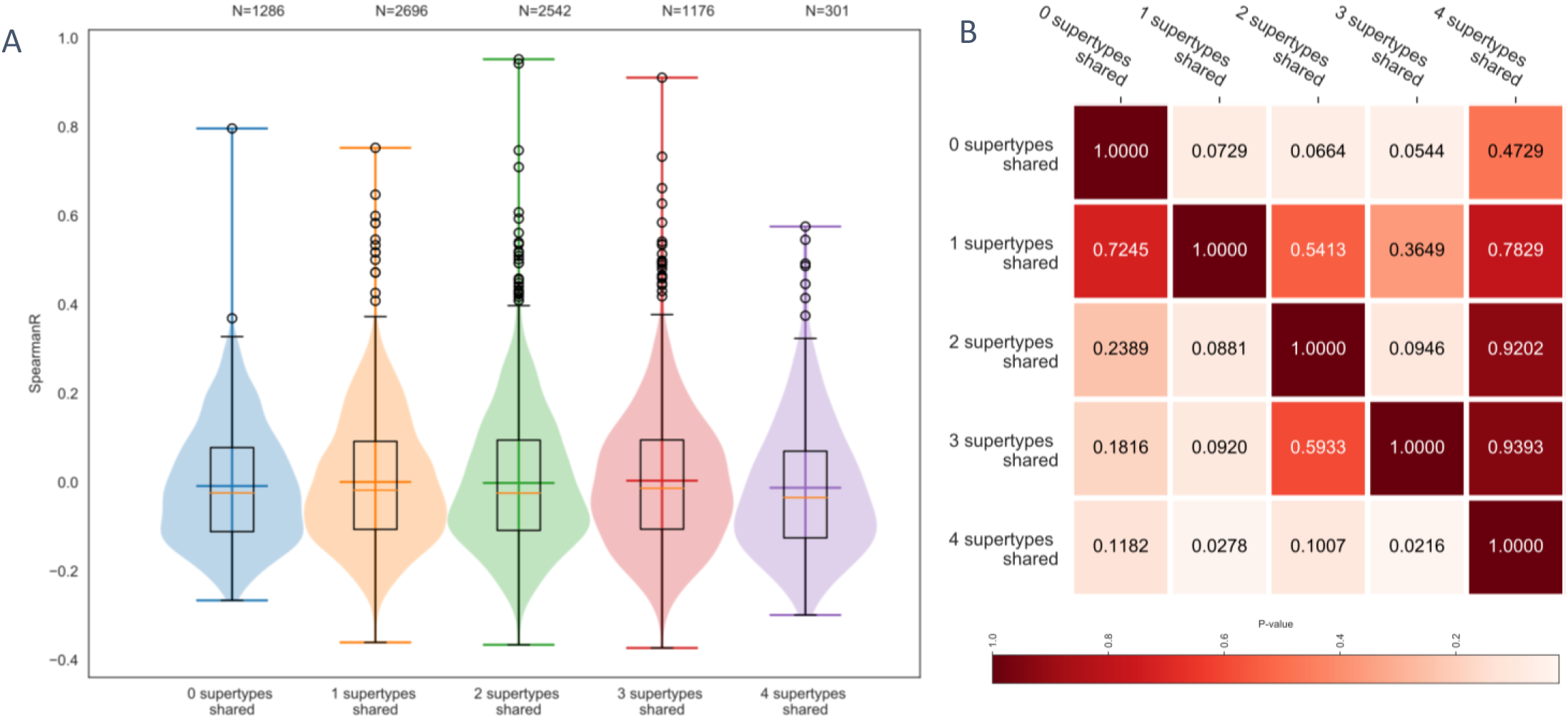
Microbiome beta diversity for samples with shared HLA class I supertypes. Spearman Rank correlation for sample pairs with 0 (blue), 1 (orange), 2 (green), 3 (red), or 4 (purple) HLA A and B supertypes shared. Heatmap on the right displays the P-values for the two sample Kolmogorov-Smirnov test under the null hypothesis; item on the y-axis is drawn from an equal or smaller distribution than the item on the x-axis.

